# Kappa opioid receptors delay sucrose self-administration by modulating dopamine ramps before operant responding

**DOI:** 10.1101/2025.10.23.684229

**Authors:** Conner W Wallace, Emanuel F Lopes, Katherine M Holleran, Sara R Jones

## Abstract

Chronic consumption of drugs of abuse increases sensitivity of the dynorphin/kappa opioid receptor (KOR) stress system, which reduces striatal dopamine signaling and promotes drug taking. However, there is mixed evidence on how KOR-modulation of dopamine affects operant behavior in the absence of addiction. To address this gap, we injected dLight1.2 in the nucleus accumbens of male and female C57BL/6J mice and trained them to self-administer sucrose. Mice were injected with the KOR agonist U50,488H, with or without the KOR antagonist aticaprant, prior to each session. U50,488H did not affect dopamine responses during the lever press and did not alter total presses or consumption. The primary effects of KOR activation were to delay initiation of lever pressing, and to diminish the rapid ramp-up in dopamine seen before lever presses. The first lever press of each session was preceded by the largest dopamine ramp; however, ramp size did not correlate with latency under any condition. We further tested whether KOR-modulation of dopamine ramps affects behavior throughout the session. We found that the total amount of dopamine activity 10 seconds before the lever press predicted when the next lever press would occur, specifically after U50,488H injection. When the phasic peaks were subtracted from this to quantify a slower dopamine ramp, time to the next lever press could be predicted and this effect was strongest after U50,488H injection. As U50,488H reduces basal dopamine signaling, we propose that dopamine ramps prior to behavior have a larger signal-to-noise ratio during KOR activation. This relatively greater increase in dopamine from baseline could increase the salience of the decision to perform reward-associated actions, which may play a role in habitual behaviors like drug taking.

## 1. Introduction

The dynorphin/kappa opioid receptor (KOR) system responds to stressful stimuli in part by exerting control over striatal dopamine (DA) neurotransmission ^1,2^. This is accomplished by locally downregulating extracellular DA concentrations via presynaptic KORs in a negative feedback loop ^3^. Upregulated KOR sensitivity due to certain stress exposures, such as early life isolation stress ^4^ or chronic forced swim stress (FSS) ^5^, induces a hypodopaminergic environment in the NAc that potentiates drug and alcohol taking. A common neurobiological mechanism promoting the reward potency of multiple classes of drugs of abuse, including alcohol ^6–8^, stimulants ^9–11^, and opioids ^12–14^, is the ability to increase extracellular DA in the nucleus accumbens (NAc) core. Consequently, overuse consistently increases the expression or sensitization of dynorphin and KORs with alcohol ^15^, stimulants ^16^, and opioids ^17^. Heightened activity of the KOR system during stress associated with drug withdrawal reduces basal DA concentrations ^18^, whereas withdrawal symptoms are alleviated and DA concentrations restored by antagonizing KORs ^16,19^. Thus, the dynorphin/KOR system is critical in controlling behavioral responses to stress and development of substance use disorders by modulating DA.

Research on the behavioral outcomes of KOR activation is less consistent in naïve or unstressed animals. Indeed, in non-dependent or unexperienced animals, acute application of both KOR agonists and antagonists have been shown to either increase or decrease the intake or rewarding properties of drugs of abuse ^2,15,20–29^. Activation of KORs in the striatum can also either promote or inhibit liking versus wanting responses for food, depending on the subregion of the NAc that is targeted ^30^, and KOR agonist modulation of sucrose intake depends on the concentration of sucrose ^31^. It has also been shown that KOR activation can reversibly decrease motivation to obtain sucrose under progressive ratio (PR) schedules of reinforcement ^32,33^, or either to have no effect on sucrose self-administration (SA) ^34^ or to slightly reduce sucrose SA ^33^ in studies that both used a fixed-ratio 5 (FR5) schedule. While application of a KOR antagonist was shown not to affect total responding for sucrose on an FR1 schedule, blocking KORs promoted faster learning of the task ^35^. There are at least two underlying theories that could explain either an increase or decrease in motivated behavior due to KOR signaling in naïve rodents. It is possible that KOR signaling either 1) inhibits motivational drive that is normally invigorated by DA, at least in drug-naïve states ^22,36,37^, or 2) promotes responding/intake of rewards to increase DA and offsets the effects of KOR activation in drug-dependent states ^5,20–22,38^. Due to the discrepancies in the literature, we aimed to test whether the KOR agonist U50,488H (U50) would increase or decrease sucrose SA behavior, and, more specifically, whether observed behavioral differences were associated with changes in real-time DA signaling, as measured by the photosensor dLight, in stress- and drug-naïve C57BL/6J mice.

The theory of reward prediction error (RPE) is well recognized in the DA field, where rapid phasic DA responses in the NAc core signal the difference between a predicted reward and the reward itself, and shift from an initially unconditioned stimulus shift to the earliest cue predicting that reward (i.e., the conditioned stimulus) ^39–42^. More recently, a different mode of DA signaling in learning and motivated behavior has emerged, which may ^43,44^ or may not ^45^ fit within the RPE framework. This is the phenomenon known as ‘DA ramps’, which may be defined as either a slow or rapid ramp-up in extracellular DA during conditioned responding that peaks at task completion or receipt of reward ^46–48^. This is distinct from classic “phasic” release events, which involve rapid release and clearance back to baseline tied to predictive cues and unexpected rewards. Many contextual modulators of DA ramps have been investigated during operant or other trained behaviors ^37,46–53^, and DA ramp activity can be seen prior to alcohol ^54^ and cocaine ^55^ SA. It has also been shown that DA ramps are larger when animals enter a heroin-paired chamber ^56^. However, these papers have quantified behavior-related effects of anticipatory signaling before a reward or a previously paired reward-associated stimulus. It is unknown whether DA ramps can be modulated directly by pharmacology, and whether KORs might control motivated behavior through modulation of pre-behavior ramps or post-learning phasic signals.

While it has been shown that a KOR antagonist promoted acquisition of both positive- and negative-valence cue association learning ^35^, and KOR agonists have been shown to either promote ^57^ or inhibit ^58^ learning and memory formation, literature on how KOR-modulation of real-time DA signaling affects operant behavior is lacking.

Given that KOR signaling reduces phasic DA signaling in the NAc by reducing phasic firing from ventral tegmental area (VTA) NAc-projecting DA neurons ^59^ and by inhibiting release locally via presynaptic receptors ^15,22,60–62^, we hypothesized that phasic-like DA signals that occurred at the time of the lever press would be diminished by U50, and that this would subsequently lead to a reduction in overall lever pressing behavior and consumption throughout the session. KOR activation also leads to a reduction in extracellular, tonic DA concentrations through a combination of the above mechanisms with increased uptake rate ^63–66^. As slow DA ramps have been shown to relate to anticipation of rewards and motivation to obtain them ^37,47,51,67^, and to track expected reward values ^52^, we predicted that slow, tonic-like DA ramps would be largest prior to the first lever press, related to the initiation of operant responding within each session, and further that a reduction in DA ramps by KOR activation would delay latency to initiate. To our surprise, there were no effects of KOR pharmacology on phasic DA responses after the lever press or on the overall amount of behavioral responding.

However, as expected, the first lever press of each session was preceded by the largest DA ramp, though this was true only for rapid (within 0.25 seconds) and not slower (10-second) DA ramps, and this rapid DA ramp was diminished after the first press prior to subsequent presses. Furthermore, KOR activation reduced the size of DA ramps overall. However, the size of the rapid DA ramp did not correlate with the time to lever press occurred from the time sucrose became available. By looking over a longer timeframe, slower DA ramps that occurred over a 10-second window up to the press negatively correlated with time to the next press, with the strongest relationship found during U50 sessions. Overall, this paper discusses how DA ramps relate to both initiation and continuation of operant behavior, with KOR activation modulating this relationship to affect the pattern of responding.

## 2. Experimental Design and Methods

### 2.1. Animals

Animal protocols were reviewed and approved by the Wake Forest University Institutional Animal Care and Use Committee. Seven-week-old C57BL/6J mice (N = 32: n = 15 males, n = 17 females; Jackson Laboratory, Bar Harbor, ME) were allowed to acclimate in our colony room for one week housed on a 12:12 light/dark cycle (lights on: 0600, lights off: 1800). Animals received water and standard rodent chow *ad libitum*. Mice were approximately 12 weeks old at the start of behavioral testing. Mice were excluded from final photometry analyses if they met at least one of the following criteria: lack of dLight signals/GFP fluorescence (n = 5 males, 3 females), incorrect cannula placement (n = 1 male), loss of headcap during training (n = 1 male, 1 female), or failure to acquire the sucrose lever pressing task (n = 4 males, 8 females). These criteria were not mutually exclusive (e.g., a mouse both failed to acquire the task and had no GFP expression upon histology). The final dataset consisted of N = 15 (9 males, 6 females) for behavioral comparisons, and N = 13 (7 males, 6 females) for DA comparisons and dLight-behavior correlations.

### 2.2. Stereotactic surgery

Mice were induced under general anesthesia using isoflurane and given meloxicam (5 mg/kg) before surgery. The surgical site was prepared and mice were mounted in a computerized stereotactic frame (Neurostar; Tübingen, Baden-Württemberg, Germany). The dLight virus (700 nL; AAV5-hSyn-dLight 1.2; Addgene, Watertown, MA; Catalogue No.: 111068-AAV5) was injected unilaterally into the NAc core (AP: +1.1 mm, ML: ± 1.3 mm, DV: +4.55 mm; from Bregma). Left versus right unilateral injection sites were counterbalanced across surgical cohorts. Next, a fiber optic cannula (Doric Lenses; Québec, Canada; Part No.: MFC_MF2.5_B280-4401-4.7) was lowered 0.10mm above the injection site (AP: +1.1 mm, ML: ± 1.3 mm, DV: +4.45 mm from Bregma). Adhesive luting cement (C&B-Metabond; Parkell; Edgewood, NY; Part No.: S380) was applied to lock the cannula in place followed by black dental cement (Lang Dental; Wheeling, IL; Part No.: 1530 & 1404) to close the surgical site. Neo-Predef with Tetracaine antibiotic powder (Zoetis; Parsipanny, NJ) was applied. Mice were monitored after surgery and administered meloxicam for ≥2 days as needed for analgesia.

### 2.3. dLight fiber photometry acquisition

Photometry recordings were taken using Fiber Photometry Systems from Tucker Davis Technologies (TDT; RZ5P; Synapse; Alachua, FL) with supporting connections purchased from Doric Lenses (Québec, Canada). Two excitation wavelengths, 405 nm (isosbestic control signal) and 465 nm (dLight1.2-dependent signal), were emitted from LEDs (CLED405, CLED465, Doric Lenses) and controlled by a multi-channel programmable LED driver (LEDD_4, Doric Lenses). These light signals were channeled into 0.57NA, 400μm core low-autofluorescence patch cords using a Doric 4-Port Fluorescence Mini Cube (FMC4, Doric Lenses), which were pre-bleached in-house for ≥4 hours on each channel. Light intensities were maintained at the tip of the patch cord at 15 μW for the 405 nm light and 15-30 μW (established during home cage testing, see **Section 2.4**) for the 465 nm light across experimental sessions. Fluorescence signals were detected using a Visible Femtowatt Photoreceiver (Newport Corporation; Irvine, CA; Model 2151). Synapse was used to modulate excitation lights (405 nm: 210 Hz, 465 nm: 330 Hz) and to measure demodulated and low-pass filtered (6 Hz) transduced fluorescence signals in real time via the RZ5P. Time-locked behavioral events were recorded in real time using Synapse, including manual experimenter-applied stimuli during home cage experiments as well as automatically generated TTL pulses from Med-PC IV (Med Associates, Inc., Fairfax, VT) via the RZ5P processor.

### 2.4. Home cage testing for dLight signals

Approximately four weeks postoperatively, mice were tested in their home cage for dLight signals. They were moved to the behavioral room and allowed to acclimate for at least 30 minutes. Following acclimation, a fiber optic patch cord was attached to their cannula via a ceramic sleeve (Precision Fiber Products; Milpitas, CA), and mice were placed back in the home cage. During this time, intensity of the 465 nm light was modulated starting at 15 μW and gradually increased to 30 μW until clear signals were seen. Mice were exposed to stimuli, including delivery of sucrose pellets to the home cage, tail pinch, air puff with a transfer pipette, and a stronger air puff with compressed air. This was to confirm presence of dLight signals prior to behavioral training. Mice deemed to have acceptable signals started sucrose SA training ≥2 days later.

### 2.5. Sucrose self-administration (SA) task

Mice were trained to self-administer 20 mg sucrose pellets (TestDiet, Part No.: 1811555, 5TUT) in classic-sized mouse operant boxes (Med Associates, Inc., Fairfax, VT, Part No. ENV-307A). For all sessions (training and pharmacological testing), when the session started, the house light and fan turned on, levers extended, and the cue lights over each lever flashed. Mice underwent no more than one training session per day and no more than two pharmacological challenges per week after acquisition. Each session lasted one hour. If the maximal responses allowed (60 pellets delivered) were reached, the boxes would turn off, but mice were allowed to remain in the chamber for the full hour. For all sessions, mice responded on a FR1 schedule wherein each press on either left or right lever resulted in immediate presentation of the cue light and delivery of one sucrose pellet. There was no inactive lever. For the initial phase of training, mice were non-tethered and able to respond on the levers, but also received free, random deliveries of sucrose pellets in conjunction with cue light presentation.

Random delivery was determined by sampling from an 11-item Fleshler-Hoffman distribution ^68^ based on an average of 240 seconds for the variable interval schedule. After non-tethered training, mice were tethered for all sessions via the fiber optic patch cord for dLight recordings, and they no longer received free pellets for the remainder of the study. Acquisition criteria were: (1) ≥10 responses during non-tethered sessions followed by tethered responding of either (2a) ≥50% of non-tethered responding over two consecutive sessions or (2b) ≥100% of non-tethered responding in one session.

The dLight recording started approximately 1 minute prior to the start of each tethered behavioral session, and automatic TTL pulses were sent from Med Associates to Synapse to time-lock the experiment start and lever presses. dLight signals were recorded for the first 20 minutes of each session then stopped, to prevent photobleaching from obscuring effects over repeated saline and drug trials later in the study.

### 2.6. Pharmacology

Following acquisition, mice underwent two sessions weekly, preceded either by a saline control or drug injection(s). The pharmacological challenges included U50,488H (U50; 5 mg/kg; i.p.) given either alone or preceded by aticaprant (ATIC; 10 mg/kg; s.c.). Saline or U50 were given 30 minutes before the start of each session, and ATIC was given 1 hour before the start of each session. Mice were placed in their home cage after ATIC, and they were tethered and placed in the boxes immediately after saline or U50 injection, giving 30 minutes of acclimation inside the boxes before each session.

### 2.7. Drugs

Aticaprant was purchased from MedChemExpress (Product: HY-101718, CAS: 1174130-61-0) and mixed in 5% DMSO / 5% Cremophor / 90% saline at 2 mg/mL. U50,488H (RTI Log No.: 13692-54B) was generously provided by the NIH and mixed in saline at 1 mg/mL.

### 2.8. dLight fiber photometry analysis

Raw data were extracted from Synapse into a format readable by MATLAB (v. R2023b) using code from TDT. A custom MATLAB pipeline was used ^69^, which was edited to analyze time-locked behavioral events according to the timing of interest.

Briefly, the raw isosbestic control (405 nm) channel was subtracted from the experimental (465 nm) signal and transformed into %ΔF/F values. The dLight responses to session start (house light & fan turn on, cue lights flash, levers extend) and lever presses (immediate presentation of cue light and sucrose pellet delivery) were analyzed by setting a local baseline for each event. A window of time prior to each event (either −3 to −1 seconds or −10 to −8 seconds, depending on analysis) was set to a value of z-score = 0 by taking the average and variability of the %ΔF/F trace during that window. The entire z-scored trace for each event (either −3 to +3 seconds or −10 to +10 seconds, depending on analysis) shows a change in signal-to-noise ratio based on this initial window set to z = 0. The z-scored traces for each event were then averaged per animal and shown, or specific periods of time within each trace were analyzed for statistics. For the analysis that baselined −10 to −8 seconds before the lever press, we defined lever pressing bouts and only included the first press in each bout, to ensure that no presses occurred during the baseline period of the next bout. A bout was defined as a press (or series of presses) that occurred with no subsequent presses within 10 seconds, or, in other words, a 10-second break between the last press of a bout and the start of the next bout.

### 2.9. Statistical analysis

Statistical analysis was conducted via GraphPad Prism (v. 10.0.2.). Behavioral results were analyzed for lever presses, pellets consumed, latency to initiate, and inter-press interval either by averaging all values for sessions within a drug type within animal and comparing animal averages, or by comparing session average values. These were compared using two-way ANOVA with drug and sex as independent variables. To conduct statistical tests on behavior-linked dLight data, the area under the curve (AUC) of the z-scored dLight signal prior to or after each TTL pulse were taken (e.g., 0.25 to 0 seconds before the lever press, 0 to 2 seconds after the lever press, or 10 to 0 seconds before the lever press). We also compared the amplitude of DA response when the lever press occurred and the peak z-score achieved after the press. The width of responses after the response was calculated as the time it took for the signal to rise from where it started at t=0 then fall back to that same z-score value. These values were compared as animal averages using one-way ANOVA, as there were no sex effects. Correlation analysis was run between the AUC of DA ramps and the time within the session that a lever press occurred. Final, multivariable linear regression was run to predict either the size of DA ramps (either from 0.25 seconds before or 10 seconds before the press) using behavioral variables associated with each lever press, or to predict when the next lever press would occur based on Tukey’s and Šídák’s multiple comparisons tests were run post-hoc to assess effects of pharmacology and sex.

## 3. Results

### 3.1. A KOR agonist delayed task initiation after sucrose availability

Mice were trained to lever press on an FR1 schedule to obtain sucrose pellets during dLight recordings. After acquisition, animals received injections of the KOR agonist U50,488H (U50; 5 mg/kg; i.p.) or a saline (i.p.) 30 minutes prior to sessions. On some of the U50 sessions, animals received an injection of the KOR antagonist aticaprant (ATIC; 10 mg/kg; s.c.) 30 minutes prior to U50 or saline. **Figure 1** shows the effects of KOR pharmacology on the three primary behavioral metrics measured using values for all sessions in all animals, which were normalized within each animal to the average of its saline sessions. There were no effects of drug or sex on the total number of lever presses (**Figure 1A**) or pellets consumed (**Figure 1B**) when expressed as a percent of the saline session averages within each animal. However, one-way ANOVA revealed an effect of drug (F_(2,125)_ = 26.24; p<0.0001) and of sex (F_(1,125)_ = 3.969; p=0.0485) on latency to initiate lever pressing after session start. Post-hoc tests showed that U50 increased latency, which was blocked by ATIC, in males (saline: 100.0 ± 10.54%; U50: 745.5 ± 155.8%; ATIC + U50: 178.6 ± 48.10%; p<0.0001, p<0.0001) and females (saline: 100.0 ± 11.35%; U50: 448.1 ± 175.8%; ATIC + U50: 74.46 ± 25.46%; p<0.01, p=0.0264). Given that values were normalized to each animal’s saline trials, the sex effect likely stemmed from a lower effect of U50 to increase latency in females compared to males (p=0.0411). These results suggest that KOR activation modifies the latency to initiate the self-administration task without altering the subsequent behavioral architecture.

**Figure 1.**
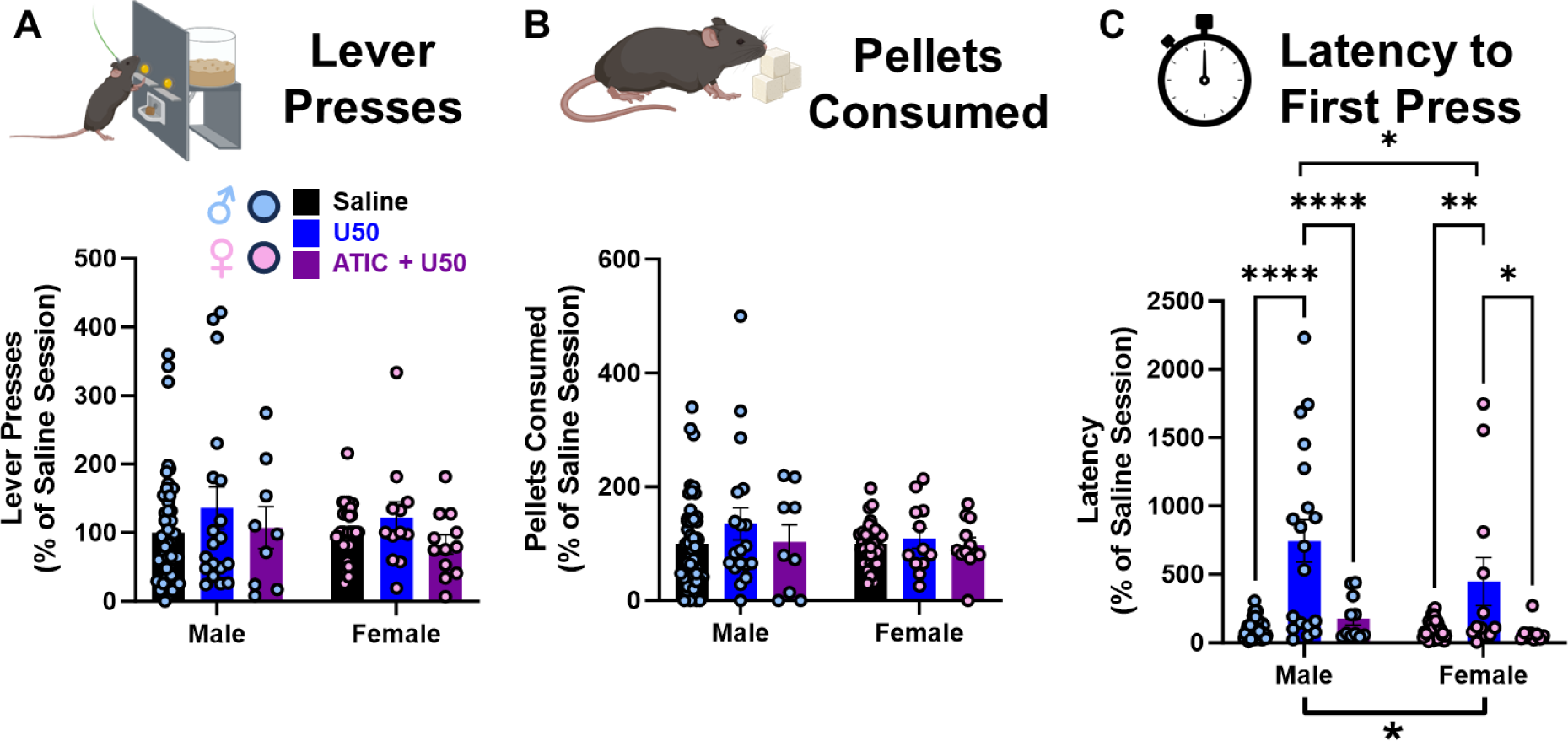
KOR activation modifies latency to first press. Mice were trained to self-administer 20 mg sucrose pellets on an FR1 schedule. The number of lever presses **(A)** and number of pellets consumed **(B)** are unchanged by KOR agonism or antagonism. KOR activation significantly increases latency to first press, and this increase is partially blocked by the KOR antagonist ATIC **(C).**

The raw lever pressing data for all sessions is shown between animals (white and gray highlighting) and drug conditions (black – saline; blue – U50; purple – ATIC + U50) in **Figure S1** for males (**Figure S1A**) and females (**Figure S1B**). The red stars indicate sessions when animals reached the maximum response limit (60 presses), which only occurred in females and was maintained throughout drug conditions. Similar to the normalized data (**Figure 1**), there were no effects of drug on total lever presses (**Figure S1C**) or pellets consumed (**Figure S1D**) in the raw data expressed as an average value per animal. However, females pressed the lever more times overall (F_(1,13)_ = 25.52; p<0.001) and consumed more pellets (F_(1,13)_ = 12.97; p<0.01) compared to males. Regarding raw latency values (**Figure S1E**), there was an effect of drug (F_(2,23)_ = 7.922; p<0.01) but no effect of sex. Post-hoc tests showed that U50 increased latency, which was blocked by ATIC, in males (saline: 165.5 ± 23.50 s; U50: 914.3 ± 233.8 s; ATIC + U50: 282.5 ± 64.12; p<0.001, p<0.01). In females, while trends towards greater latency in the U50 session and lower latency in the ATIC + U50 sessions was visually evident, this was not significant.

Overall, females pressed the lever more and consumed more sucrose compared to males, but there were no effects of KOR pharmacology in either sex on these variables. The major effect of U50 on overall behavior throughout sessions was to delay the latency to initiate lever pressing.

### 3.2. DA ramps that precede a lever press for sucrose are diminished by a KOR agonist

When each operant session started, mice were cued to the availability of sucrose by the house and fan lights turning on, cue lights flashing, and the levers extending.

After this point, mice could decide when to start pressing and how often, with no restrictions other than a maximum allowed amount of 60 presses/pellets per session. When mice pressed the lever, the cue lights briefly illuminated and a sucrose pellet was immediately delivered (**Figure 2A**). A representative DA response to the lever press is shown in **Figure 2A**. In the representative response, DA steadily increases prior to the press (a DA ramp), followed by a phasic-like signal when the cue lights flash and the pellet is delivered. The average DA responses within this 6-second window were taken for all lever presses within a given session type and within each animal. The average of these responses per animal were then shown to give an overall comparison of drug effects on DA responses to lever presses. These are shown as traces comparing saline versus U50 sessions (**Figure 2B**), U50 versus ATIC + U50 sessions (**Figure 2C**), and saline versus ATIC + U50 sessions (**Figure 2D**).

**Figure 2.**
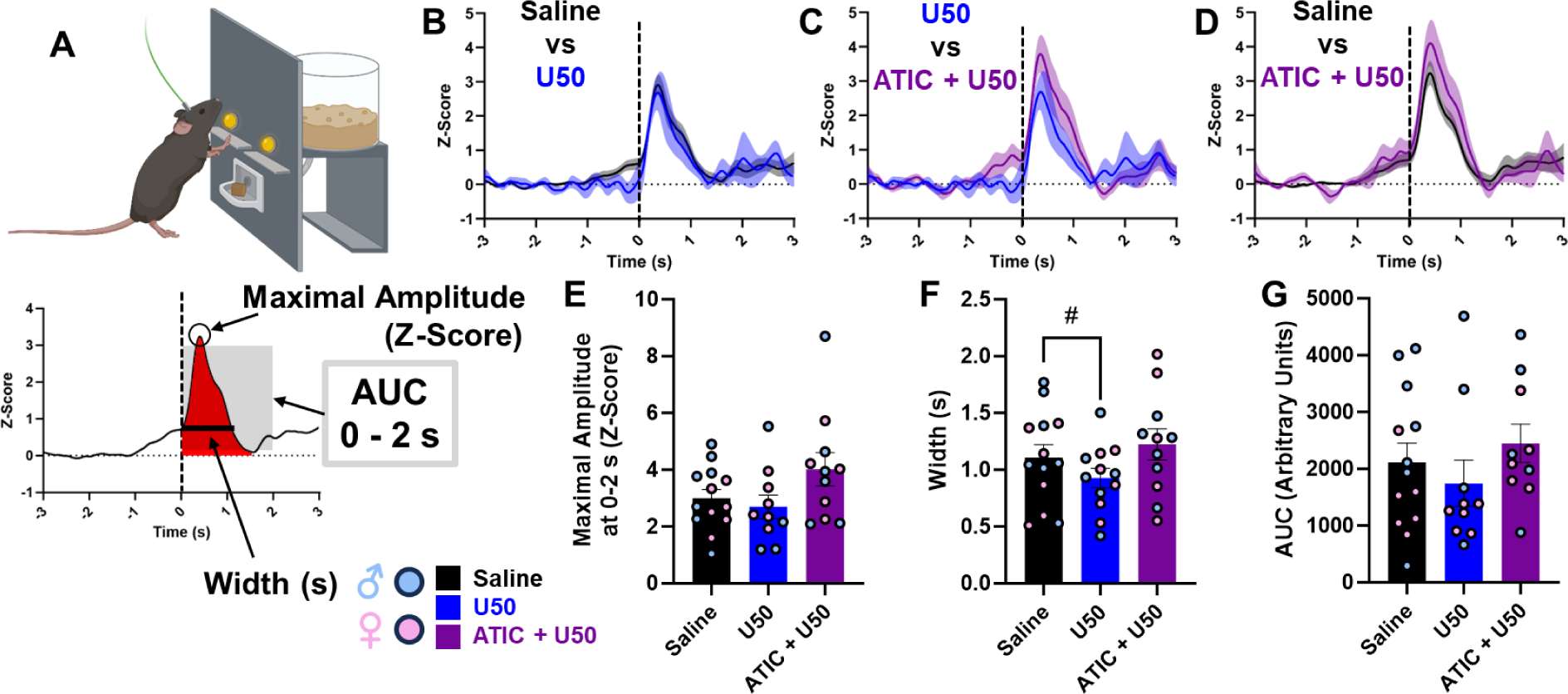
KOR modulation did not significantly affect dLight DA responses following the lever press. **(A)** Schematic highlighting a representative DA response to the lever press, along with parameters of the response being assessed (Maximal amplitude, width, AUC 0-2s). Average DA responses 6 seconds surrounding the lever press in saline vs U50 sessions **(B)**, U50 versus ATIC + U50 sessions **(C)**, and saline versus ATIC + U50 sessions **(D)**. **(E)** Maximal amplitude of DA responses in saline, U50, and ATIC + U50 sessions. **(F)** Width of DA responses in saline, U50, and ATIC + U50 sessions. **(G)** AUC of DA responses in saline, U50, and ATIC + U50 sessions. No significant effect of drug was identified as DA responses following the lever press.

The average DA responses per animal were next quantified. We first looked at the maximal amplitude of dLight response after the lever was pressed. There was a trend towards an effect of drug on amplitude (F_(1.361,12.93)_ = 3.635; p=0.0693), but this was not significant (**Figure 2E**). We next quantified the width of each dLight response, i.e., the time it took for the z-score to rise from t=0s and fall back to the z-score value at that t=0. There was a trend for a drug effect on width (F_(1.406,14.76)_ = 3.257; p=0.0807) with post-hoc tests showing a trend for U50 to reduce width (saline: 1.107 ± 0.1143 s; U50: 0.9287 ± 0.08535 s; p=0.0956), but neither of these effects reached significance (**Figure 2F**). Finally, the area under the curve (AUC) of dLight signal from 0 to 2 seconds after the press was taken, but there was not a significant drug effect (F_(1.309,11.79)_ = 0.8415; p=0.4085; **Figure 2G**). Overall, KOR modulation did not significantly affect DA responses following the lever press.

We next quantified whether U50 affected DA ramps leading up to the lever press, and we focused on a time window preceding the lever press (−2 to 0 seconds, **Figure 3A**). It is visually evident that DA ramps were smaller in the U50 compared to saline condition (**Figure 3B**) and compared to the ATIC + U50 condition (**Figure 3C**), and DA ramps looked similar between saline and ATIC + U50 conditions (**Figure 3D).** A one-way ANOVA showed a significant effect of drug on the amplitude of dLight signal at time t = 0s (F_(1.754,15.78)_ = 5.005; p=0.0240; **Figure 3E**). Post-hoc tests showed that U50 reduced the amplitude at this timepoint (saline: 0.62 ± 0.15 z-score, U50: 0.052 ± 0.1244 z-score; p=0.0321), and there was a trend for ATIC to block this effect (ATIC + U50: 0.47 ± 0.19; p=0.0885). We also quantified DA ramps by calculating the area under the curve (AUC, arbitrary units) between the lever press and 0.25 seconds before the lever press. A one-way ANOVA revealed a drug effect on DA ramp AUC (F_(1.817,16.35)_ = 5.654; p=0.0155; **Figure 3F**). Post-hoc tests showed a trend for U50 to reduce the size of this response (saline: 143.5 ± 37.3 AUC, U50: 30.9 ± 11.3 AUC; p=0.05694). Furthermore, the ATIC + U50 response was significantly higher than U50 alone (ATIC + U50: 125.0 ± 34.1; p= 0.0225). Thus, while DA responses to the lever press and sucrose delivery were unchanged, KOR activation reduced the size of preceding DA ramps.

**Figure 3.**
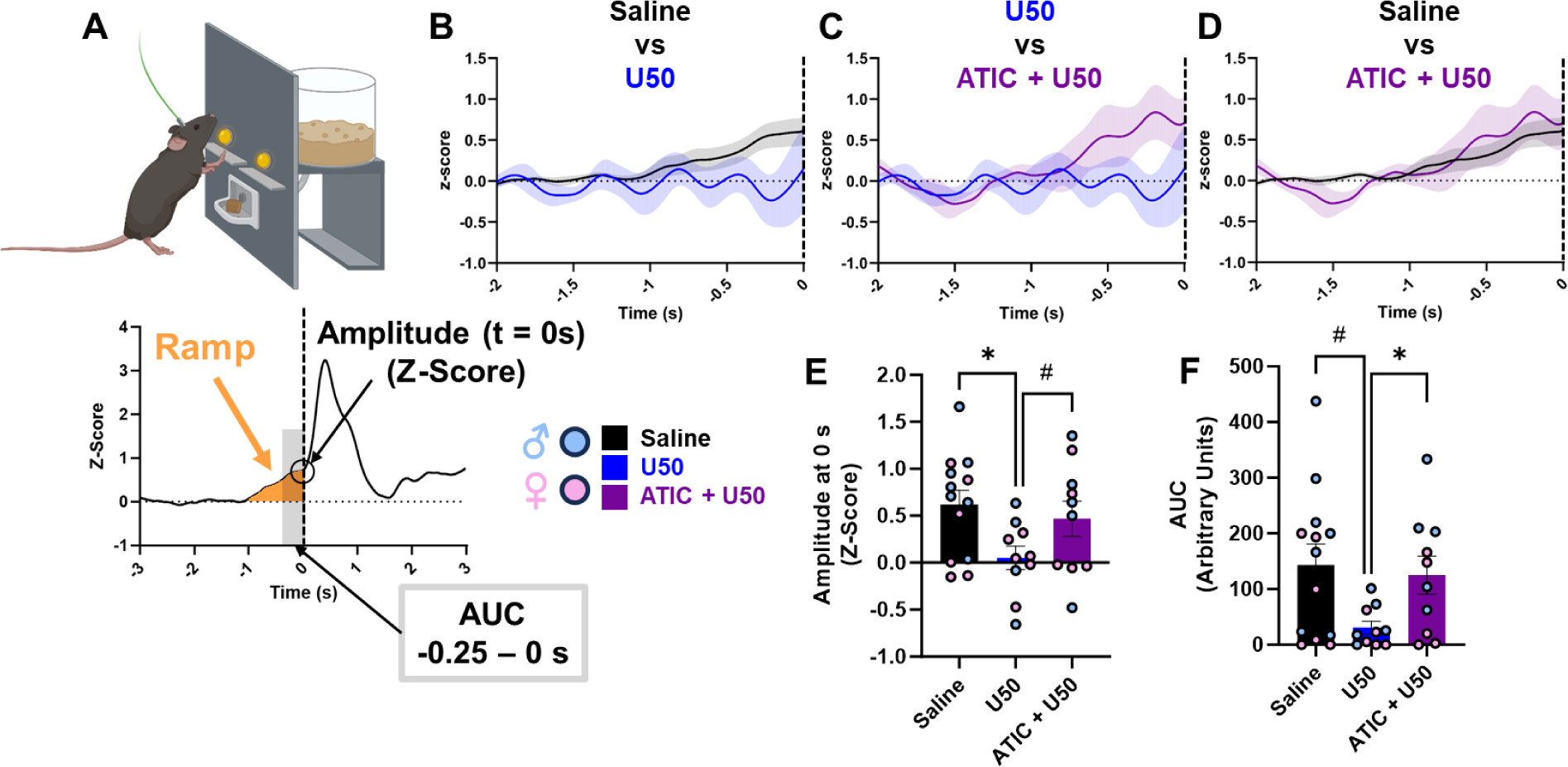
KOR modulation modifies DA ramps preceding lever presses. **(A)** Schematic highlighting a representative DA response to the lever press, along with parameters of the response being assessed (Amplitude at t=0, AUC −0.25 – 0s). Average DA responses 2 seconds before the lever press in saline vs U50 sessions **(B)**, U50 versus ATIC + U50 sessions **(C)**, and saline versus ATIC + U50 sessions **(D)**. **(E)** Amplitude of DA responses at t=0 in saline, U50, and ATIC + U50 sessions. U50 significantly reduces the amplitude of the DA ramp. **(F)** AUC (−0.25-0s) of DA responses in saline, U50, and ATIC + U50 sessions. KOR modulation trends towards a reduction in AUC, which is significantly different from the ATIC + U50 sessions.

### 3.3. Task initiation is preceded by large and rapid DA ramps, but KOR modulation of DA ramps is not related to the delay in task initiation

The primary effects of U50 shown so far were to increase the latency to initiate lever pressing and to diminish the size of DA ramps preceding a lever press.

Consequently, we next asked whether the size of DA ramps was linked to when a lever press occurred throughout the session. In **Figure 2-3**, the baseline was set from data −3 to −1 seconds prior to a lever press. However, if there was a prolonged DA ramp prior to 1 second before the press, this would have been artificially included in the baseline. For the next portion of analysis, we set a baseline of −10 to −8 seconds before the press and set the overall analysis window at −10 to 10 seconds surrounding the lever press. This allowed us to interrogate DA ramps on a longer timescale. To exclude the possibility that this longer time window might capture signals related to prior lever presses, which would interfere with baselining and z-score analysis, we defined bouts as a lever press or series of lever presses followed by a ≥10 second break. Only the first lever press in each bout was included, and several presses were consequently excluded from dLight analysis (male saline: 109 of 248; female saline: 464 of 823; male U50: 3 of 16; female U50: 63 of 91; male ATIC + U50: 11 of 27; female ATIC + U50: 63 of 101). Because all bouts were included in correlation and regression analyses, rather than animal averages, we only used the first drug session completed in each animal where lever presses occurred during the dLight recording. Every dLight trace for a lever press bout that was quantified in the remaining analyses is shown in **Figure S2** for saline (**Figure S2A**), U50 (**Figure S2B**), and ATIC + U50 (**Figure S2C**) sessions. Ramps were quantified using AUC before the first lever press in each bout for either 0.25 seconds (significantly changed by U50 in **Figure 3**) and for the entire 10 seconds period before the press. Contrary to our expectation, the size of the dopamine ramp did not correlate with when a lever press occurred within a session (session time (s)) when quantified 0.25 seconds before the ramp in saline (**Figure 4A**), U50 (**Figure 4B**), or ATIC + U50 (**Figure 4C**) conditions. Ramp size from −10 to 0 seconds also did not correlate with session time in saline (**Figure 4D**), U50 (**Figure 4E**), or ATIC+U50 (**Figure 4F**) conditions. This suggests that the U50-mediated reduction in DA ramp size does not relate to the delay in task initiation.

**Figure 4.**
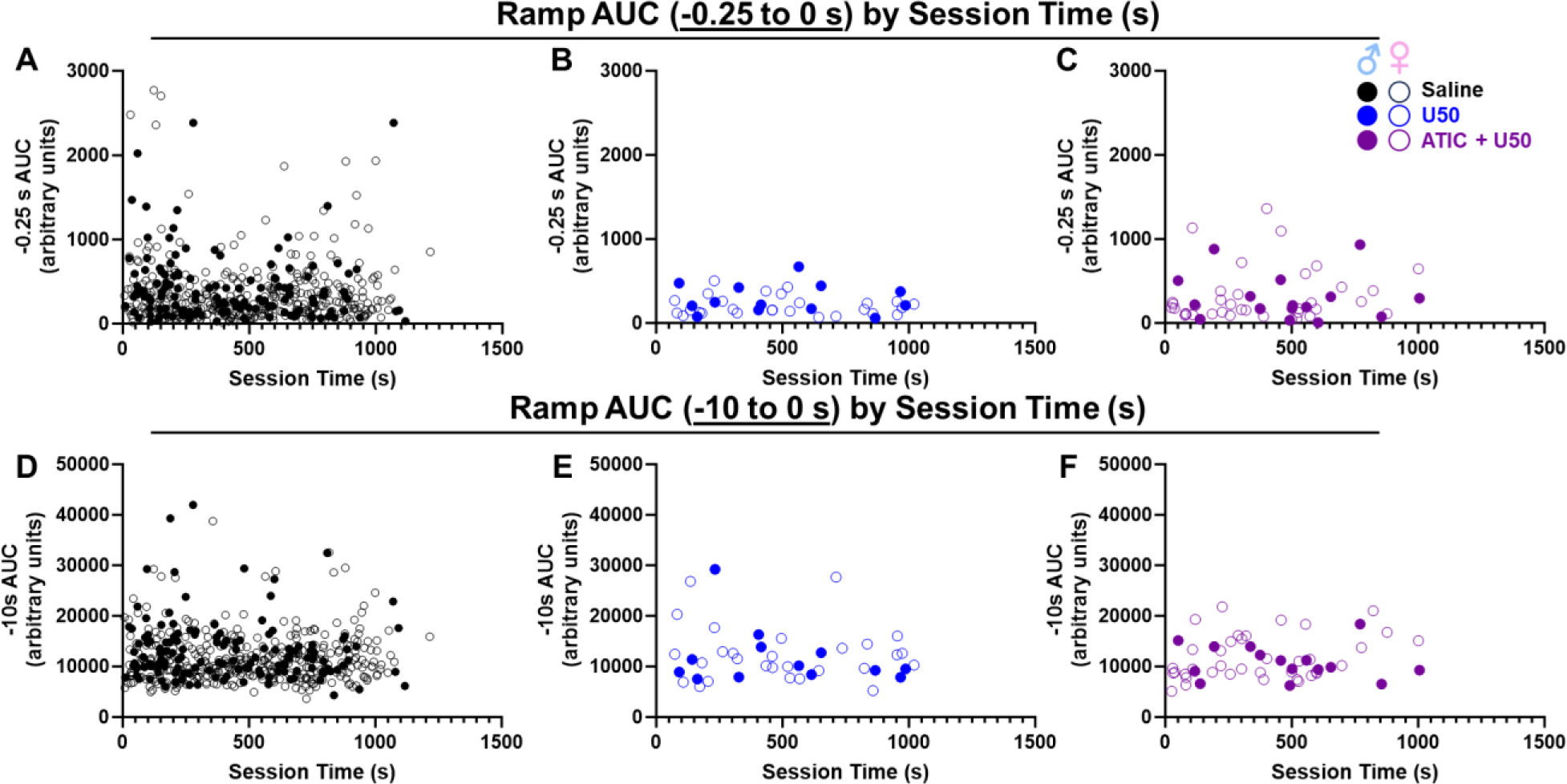
Dopamine ramp size does not correlate with lever press session time. Ramps were quantified using AUC 0.25 seconds before the first lever press in each bout and correlated with their respective session times for saline **(A)**, U50 **(B)**, and ATIC + U50 **(C)**. Similarly, ramps were quantified using AUC 10 seconds before the first lever press in each bout and correlated with their respective session times for saline **(D)**, U50 **(E)**, and ATIC + U50 **(F).** No correlation was found between AUC and session time for any drug condition or ramp duration.

Due to this apparently conflicting result, we next queried what behavioral variables might predict ramp size across conditions. We conducted a series of multilinear regression analyses to predict ramp size from the following independent variables: 1) session time, 2) bout order, 3) presses/bout, 4) whether the bout was the first in the session or not, 5) sex, and 6) drug (**Table 1**). We confirmed our prior finding and showed that session time and bout order cannot predict the size of the rapid, 0.25 second DA ramp. However, the regression found that whether the bout was the first/not first in a session predicts ramp size, i.e., the first bout in a session predicts larger ramp size (**Table 1**, **Figure S3**. Furthermore, there was a significant effect of drug, with bouts occurring during U50 sessions being preceded by smaller ramps (**Table 1**). This further confirms the findings from Figure 3, as the multivariable linear regression used (a) a ramp AUC that was analyzed using a different baseline and (b) only included the first press in each bout. This suggests that KOR-induced reduction in DA ramps prior to operant behavior is a robust finding. We ran another multivariable linear regression with the same independent variables, and none of them significantly predicted the AUC of DA activity 10 seconds prior to the bout (**Table S1**). Overall, it appears that DA ramps, specifically rapid DA ramps immediately before the press, are important for task initiation (i.e., the first lever press). While U50 reduces the size of rapid DA ramps, this is not related to the timing of task initiation.

**Table 1.**
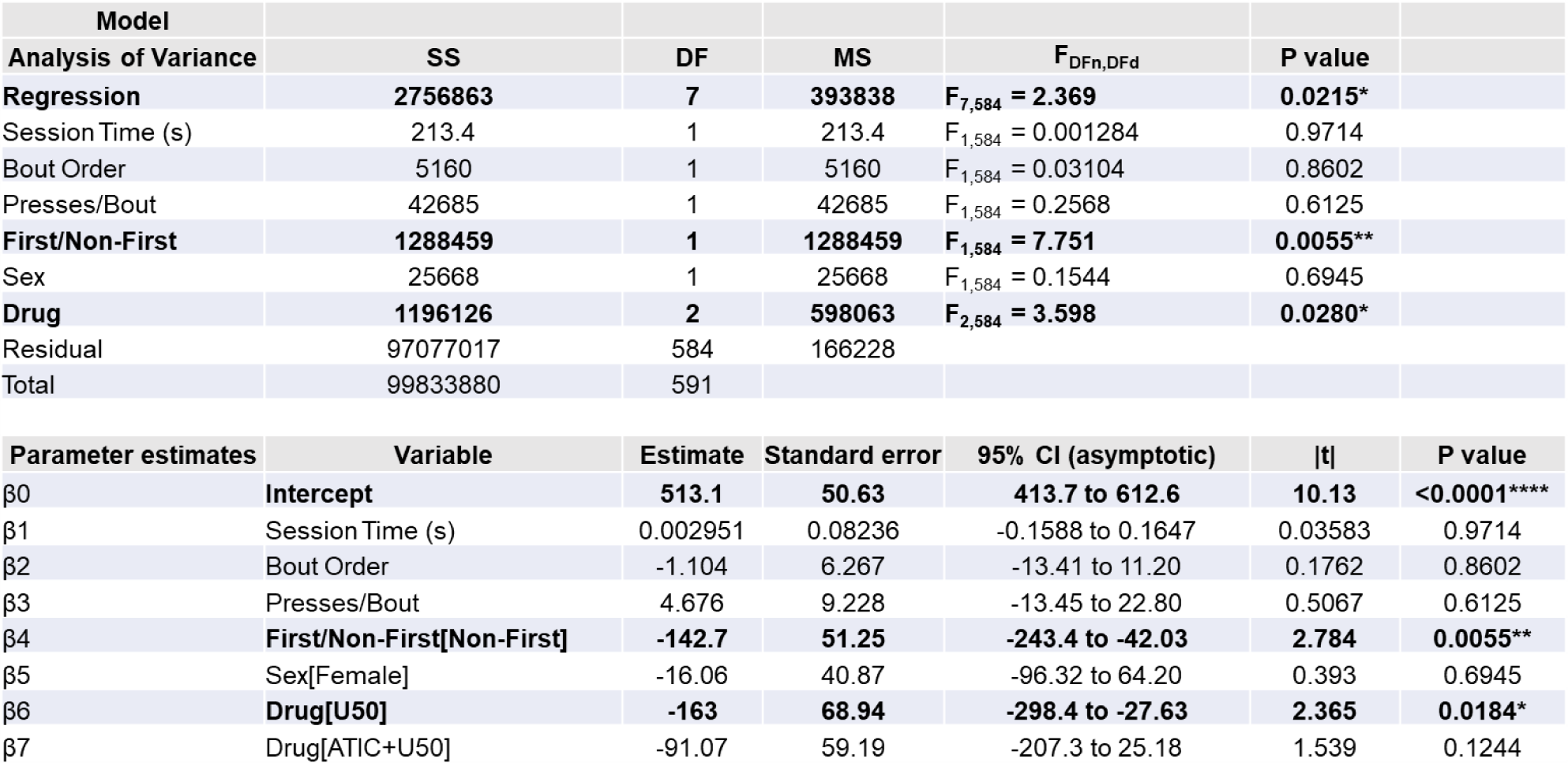
A multivariable linear regression analysis to predict the total AUC from - 0.25 seconds up to the lever press from multiple behavioral and categorical variables. The regression equation was significant, meaning the independent variables were able to predict the size of the dopamine ramp 0.25 seconds before a lever press. There was a significant drug effect, with U50 reducing the size of ramps. Further, there was a significant effect of bout order, meaning the first lever pressing bout in a session was likely to have larger ramps compared to subsequent presses.

| Model |  |  |  |  |  |
| --- | --- | --- | --- | --- | --- |
| Analysis of Variance | SS | DF | MS | F<sub>DFn,DFd</sub> | P value |
| **Regression** | **2756863** | **7** | **393838** | **F<sub>7,584</sub> = 2.369** | **0.0215\*** |
| Session Time (s) | 213.4 | 1 | 213.4 | F<sub>1,584</sub> = 0.001284 | 0.9714 |
| Bout Order | 5160 | 1 | 5160 | F<sub>1,584</sub> = 0.03104 | 0.8602 |
| Presses/Bout | 42685 | 1 | 42685 | F<sub>1,584</sub> = 0.2568 | 0.6125 |
| **First/Non-First** | **1288459** | **1** | **1288459** | **F<sub>1,584</sub> = 7.751** | **0.0055\*\*** |
| Sex | 25668 | 1 | 25668 | F<sub>1,584</sub> = 0.1544 | 0.6945 |
| **Drug** | **1196126** | **2** | **598063** | **F<sub>2,584</sub> = 3.598** | **0.0280\*** |
| Residual | 97077017 | 584 | 166228 |  |  |
| Total | 99833880 | 591 |  |  |  |

**Table 1. A multivariable linear regression analysis to predict the total AUC from - 0.25 seconds up to the lever press from multiple behavioral and categorical variables.**
| Parameter estimates | Variable | Estimate | Standard error | 95% CI (asymptotic) | \|t\| | P value |
| --- | --- | --- | --- | --- | --- | --- |
| β<sub>0</sub> | **Intercept** | **513.1** | **50.63** | **413.7 to 612.6** | **10.13** | **<0.0001\*\*\*\*** |
| β<sub>1</sub> | Session Time (s) | 0.002951 | 0.08236 | -0.1588 to 0.1647 | 0.03583 | 0.9714 |
| β<sub>2</sub> | Bout Order | -1.104 | 6.267 | -13.41 to 11.20 | 0.1762 | 0.8602 |
| β<sub>3</sub> | Presses/Bout | 4.676 | 9.228 | -13.45 to 22.80 | 0.5067 | 0.6125 |
| β<sub>4</sub> | **First/Non-First[Non-First]** | **-142.7** | **51.25** | **-243.4 to -42.03** | **2.784** | **0.0055\*\*** |
| β<sub>5</sub> | Sex[Female] | -16.06 | 40.87 | -96.32 to 64.20 | 0.393 | 0.6945 |
| β<sub>6</sub> | **Drug[U50]** | **-163** | **68.94** | **-298.4 to -27.63** | **2.365** | **0.0184\*** |
| β<sub>7</sub> | Drug[ATIC+U50] | -91.07 | 59.19 | -207.3 to 25.18 | 1.539 | 0.1244 |

### 3.4. KOR activation increases the time between bouts and strengthens the relationship between prolonged DA ramps and future behavior

We were unable to predict the size of a 10-second DA ramp from any of the behavioral variables so far. Given the importance of DA ramps in the literature of shaping behavior within sessions and over time longitudinally ^37,46,49^, we next asked a nuanced question – can the size of these prolonged DA ramps predict architecture of lever pressing bouts over a session. We quantified the length of time between the start of each bout (time to next bout (s)). Indeed, a one-way ANOVA showed a significant effect of drug on time to next bout (F_(2,445)_ = 35.73; p<0.0001; **Figure 5A**). Post-hoc tests showed that U50 increased the increased the time to next bout compared to saline (saline: 69.42 ± 2.635 s; U50: 169.8 ± 28.67 s; p<0.0001; **Figure 5A**), which was blocked by ATIC (ATIC + U50: 104.8 ± 11.32; p<0.01; **Figure 5A**). This was not a full blockade, as the ATIC + U50 condition had a larger time to next bout than saline (p<0.01; **Figure 5A**). Given the trend of the prolonged ramp and the significant effect of U50 to predict when the next bout would occur, we asked whether there was a relationship between these variables. We performed the same multivariable linear regression with two-way interaction terms, and there was a significant interaction between ramp size and U50 sessions to predict when the next bout would occur (**Table 2**, **Figure 5B).** In specifically the U50 condition, there was a negative correlation between 10-second ramp size and time to the next bout (r=-0.4033; p=0.0333; **Figure 5C**). Thus, the total amount of DA activity before a lever press bout may shape subsequent behavior, and KOR activation strengthens this relationship.

**Figure 5.**
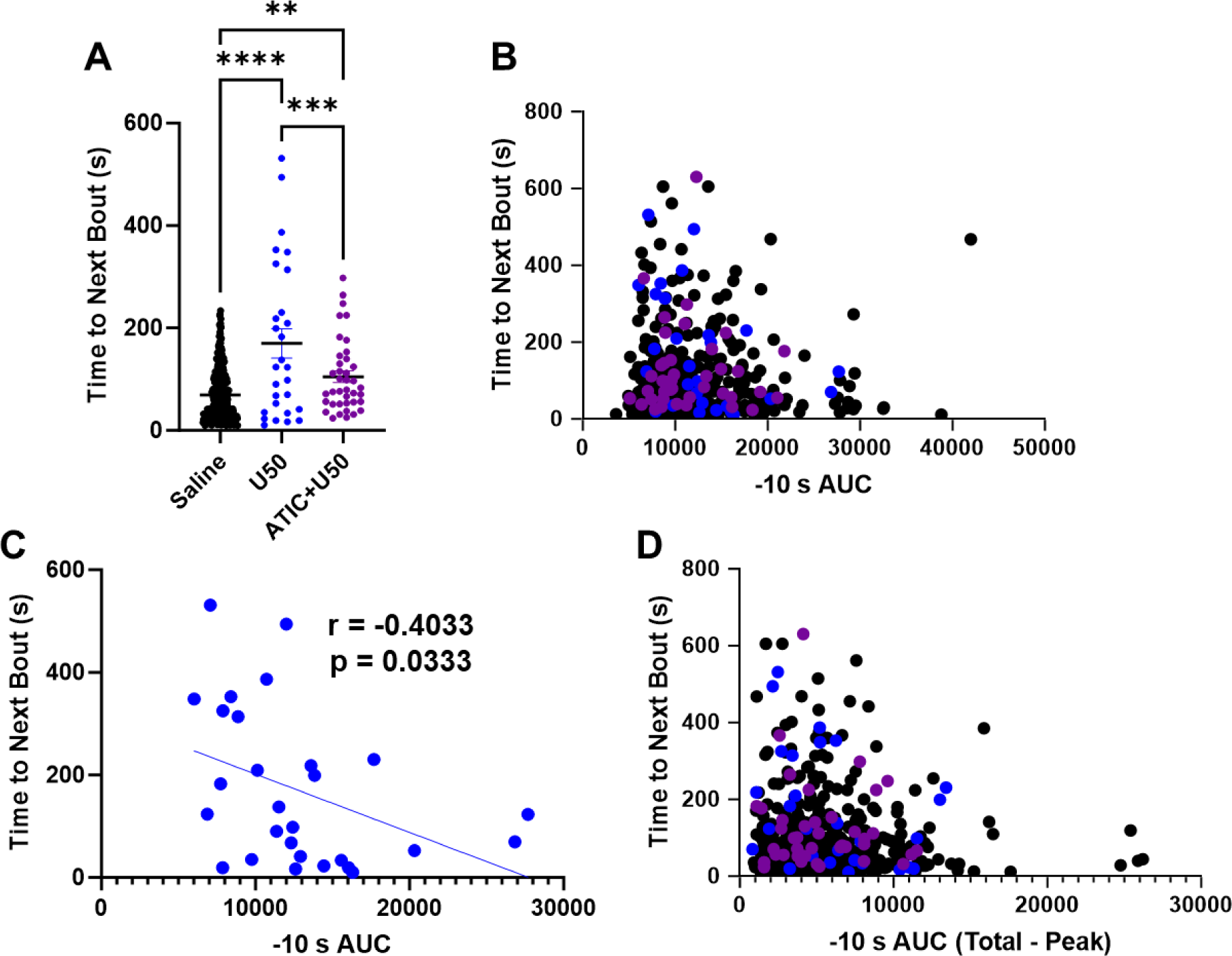
KOR activation both increases inter-bout-interval and strengthens the relationship between timing and the size of DA ramps. **(A)** The time between bouts is increased by U50, which is blocked by ATIC. **(B)** While there is not an ability overall to predict time to the next bout from the total AUC of DA activity before a given bout, this relationship is significant during **(C)** U50 sessions. When the peak AUC is subtracted from the total AUC 10 seconds before the press **(D)**, the size of this slow, tonic-like DA ramp predicts time to the next bout.

**Table 2.** A multivariable linear regression analysis to predict the time to the next bout from the total size of the AUC of DA ramps 10 seconds prior to the bout. Several independent variables were able to predict time to the next bout, including drug condition, showing that there was increased time between bouts in the U50 condition. There was a significant interaction between ramp size and the U50 drug condition, such that there was a negative correlation between ramp size and time to the next bout after U50 injection. There was a near-significant effect of sex with a negative estimate, suggesting females had less time between bouts.

| Model |  |  |  |  |  |  |
| --- | --- | --- | --- | --- | --- | --- |
| Analysis of Variance | SS | DF | MS | $F_{DFn,DFd}$ | P value | |
| Regression | 706393 | 9 | 78488 | $F_{9,479} = 7.778$ | <0.0001**** | |
| Residual | 4833462 | 479 | 10091 |  |  |  |
| Total | 5539855 | 488 |  |  |  |  |

**Table 2. A multivariable linear regression analysis to predict the time to the next bout from the total size of the AUC of DA ramps 10 seconds prior to the bout.**
| Parameter estimates | Variable | Estimate | Standard error | 95% CI (asymptotic) | \|t\| | P value |
| --- | --- | --- | --- | --- | --- | --- |
| $\beta_0$ | Intercept | 148.3 | 23.53 | 102.1 to 194.6 | 6.305 | <0.0001**** |
| $\beta_1$ | -10-0 AUC | -0.0001561 | 0.001602 | -0.003304 to 0.002992 | 0.09741 | 0.9224 |
| $\beta_2$ | Sex[Female] | -52.72 | 27.39 | -106.5 to 1.102 | 1.925 | 0.0549 |
| $\beta_3$ | Drug[U50] | 150.3 | 57.02 | 38.22 to 262.3 | 2.635 | 0.0087** |
| $\beta_4$ | Drug[ATIC+U50] | 37.65 | 54.98 | -70.38 to 145.7 | 0.6848 | 0.4938 |
| $\beta_5$ | -10-0 AUC : Sex[Female] | -0.001227 | 0.001949 | -0.005058 to 0.002603 | 0.6295 | 0.5293 |
| $\beta_6$ | -10-0 AUC : Drug[U50] | -0.01029 | 0.003845 | -0.01785 to -0.002738 | 2.677 | 0.0077** |
| $\beta_7$ | -10-0 AUC : Drug[ATIC+U50] | 0.001665 | 0.004006 | -0.006207 to 0.009537 | 0.4157 | 0.6778 |
| $\beta_8$ | Sex[Female] : Drug[U50] | 76.17 | 46.71 | -15.61 to 167.9 | 1.631 | 0.1036 |
| $\beta_9$ | Sex[Female] : Drug[ATIC+U50] | -37.76 | 38.58 | -113.6 to 38.04 | 0.9788 | 0.3281 |

As a final analysis, we wanted to assess the prolonged DA ramp 10 seconds before a lever press separate from the phasic peaks evident in **Figure S2**. While phasic peaks have been shown to occur on top of DA ramps and relate to behavioral processing along with the slower ramp ^43,47^, we wanted to know if the smaller, tonic-like increase in DA might have a separate function. We subtracted the peak AUC from the total AUC and ran the same analysis discussed immediately above. The size of the ‘total AUC’ – ‘peak AUC’ 10 seconds before the press significantly predicted when the next bout would occur overall (**Table 3**, **Figure 5D**). Again, there was a significant interaction between AUC and U50 sessions, suggesting this relationship between tonic-like ramp size and time to the next press to be strongest in the presence of U50. Of note, the size of the rapid DA ramp 0.25 seconds before a press was unable to predict when a subsequent bout would occur (**Table S2**). Overall, it is the prolonged ramp of DA 10 seconds before a lever press that predicts subsequent behavior, particularly after subtracting out the phasic events, and this relationship is strengthened by KOR activation.

**Table 3.**
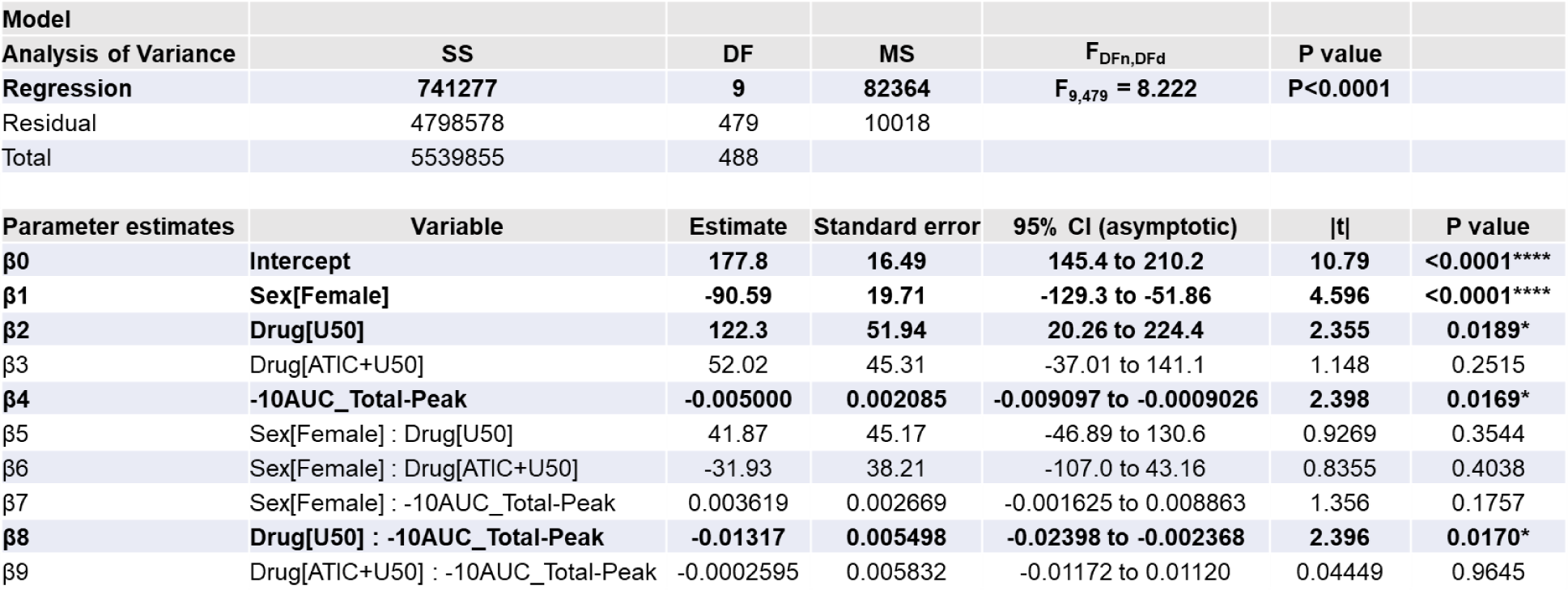
A multivariable linear regression analysis to predict the time to the next bout from slow, tonic-like DA ramps 10 seconds prior to the bout. Several independent variables were able to predict time to the next bout, most importantly here the size of DA ramps 10 seconds prior calculated as the total AUC minus the peak AUC. This was our measure of a slow, tonic-like DA ramp with the phasic-like peaks subtracted. Drug condition and sex were also significant predictors, specifically showing that U50 sessions had more time between bouts while females had less time between bouts. Finally, there was an interaction between the size of the DA ramp (total minus the peak AUC) with U50 sessions. Thus, while the slow, tonic-like DA ramp predicted when the next bout would occur overall, this relationship was, again, strongest after U50 injection.

| Model |  |  |  |  |  |  |
| --- | --- | --- | --- | --- | --- | --- |
| Analysis of Variance | SS | DF | MS | $F_{DFn,DFd}$ | P value | |
| Regression | 741277 | 9 | 82364 | $F_{9,479} = 8.222$ | $P < 0.0001$ | |
| Residual | 4798578 | 479 | 10018 |  |  |  |
| Total | 5539855 | 488 |  |  |  |  |

**Table 3. A multivariable linear regression analysis to predict the time to the next bout from slow, tonic-like DA ramps 10 seconds prior to the bout.**
| Parameter estimates | Variable | Estimate | Standard error | 95% CI (asymptotic) | \|t\| | P value |
| --- | --- | --- | --- | --- | --- | --- |
| $\beta_0$ | Intercept | 177.8 | 16.49 | 145.4 to 210.2 | 10.79 | <0.0001**** |
| $\beta_1$ | Sex[Female] | -90.59 | 19.71 | -129.3 to -51.86 | 4.596 | <0.0001**** |
| $\beta_2$ | Drug[U50] | 122.3 | 51.94 | 20.26 to 224.4 | 2.355 | 0.0189* |
| $\beta_3$ | Drug[ATIC+U50] | 52.02 | 45.31 | -37.01 to 141.1 | 1.148 | 0.2515 |
| $\beta_4$ | -10AUC_Total-Peak | -0.005000 | 0.002085 | -0.009097 to -0.0009026 | 2.398 | 0.0169* |
| $\beta_5$ | Sex[Female] : Drug[U50] | 41.87 | 45.17 | -46.89 to 130.6 | 0.9269 | 0.3544 |
| $\beta_6$ | Sex[Female] : Drug[ATIC+U50] | -31.93 | 38.21 | -107.0 to 43.16 | 0.8355 | 0.4038 |
| $\beta_7$ | Sex[Female] : -10AUC_Total-Peak | 0.003619 | 0.002669 | -0.001625 to 0.008863 | 1.356 | 0.1757 |
| $\beta_8$ | Drug[U50] : -10AUC_Total-Peak | -0.01317 | 0.005498 | -0.02398 to -0.002368 | 2.396 | 0.0170* |
| $\beta_9$ | Drug[ATIC+U50] : -10AUC_Total-Peak | -0.0002595 | 0.005832 | -0.01172 to 0.01120 | 0.04449 | 0.9645 |

## 4. Discussion

This is the first study of its kind to assess effects of KOR-targeted drugs on DA signaling in the NAc *in vivo* using fiber photometry paired with the photosensor dLight while animals were engaged in a task to obtain sucrose. The KOR agonist U50,488H (U50) significantly delayed task initiation and diminished the size of the rapid DA ramp that occurred 0.25 seconds before lever presses occurred, effects which were KOR-specific and blocked by the KOR antagonist aticaprant (ATIC). Further, the first lever press in any given session was preceded by a larger, rapid DA ramp. However, these facts were unrelated, as ramp size did not correlate with when lever presses occurred. We consequently queried whether DA ramps were related in this paradigm to future behavior, as previously shown ^49^. The rapid DA ramp immediately prior to the lever press was primarily predicted by whether a lever pressing bout was the first bout in a session, and it could not be predicted by bout order or overall timing of lever presses. Conversely, the prolonged, slow and tonic-like increase in DA that started 10 seconds before each lever press predicted when future bouts would occur, and this relationship was strengthened after U50 injection.

NAc DA has well-recognized roles in associative learning ^41^ and reward prediction error (RPE) ^70^ signaling. More recently, DA ramping, where DA activity and/or release increases as an animal approaches a reward ^71^, has emerged as novel signaling mode. DA ramping was first defined in 2013 using *in vivo* FSCV ^46^, showing that dopamine signals ramp up in the ventromedial but not dorsolateral striatum while rats navigate a maze to obtain natural rewards. In contrast to the phasic, burst-like dopamine signals expected to shift from reward to cue during learning while running a T-maze, the authors showed that DA ramped up according to spatial proximity to the reward, peaking at the time of retrieval. Since then, many studies have shown in multiple paradigms that DA ramps encode various aspects of reward, including magnitude of expected reward ^46,47,67^, external and internal processing of proximity to reward ^46,67^, anticipation of reward as cues predict its approach ^47,67^, and state uncertainty (i.e., changing probability of receiving the reward) ^51,52^. It is interesting to note that the rapid DA ramps we demonstrated immediately prior to operant behavior look similar to those reported as early as 2004 in an FR1 schedule of sucrose solution reinforcement ^48^ where cues were also presented during lever pressing.

It has also been shown that DA ramps associate with task initiation when multiple trials were given in a single session and rats had to wait for a cue before responding, and this association was strongest when rats initiated an action during successful rather than unsuccessful trials ^49^. Though our paradigm was simple, and mice only had correct/rewarded opportunities, this aligns with the fact that we showed rapid DA ramps were largest when animals initiated the first lever press of the session. After having gone several days between sessions, it may be possible that this rapid DA ramp is necessary to start lever pressing on subsequent sessions.

We show that U50 reduced the size of rapid DA ramps immediately prior to lever pressing. U50 also delayed task initiation without affecting overall behavior, suggesting that locomotion was not impaired, consistent with prior evidence using similar intermediate doses of U50 ^72,73^. Additionally, most studies have shown that the timing and size of DA ramps do not strongly correlate with movement (running speed, licking, locomotor activity, wheel turns, etc.) but rather with the anticipation of a previously conditioned reward ^46,47,67^, suggesting that DA ramps motivate initiation of a behavior but do not themselves generate the actions necessary for the behavior. Therefore, as U50 both delayed task initiation and reduced the size of DA ramps, we predicted that there would be a relationship between ramp size and either latency to the first lever press or session time when presses occurred. In other words, rather than inhibiting the ability to generate movement, we predicted that U50 diminished the size of DA ramps, thereby delaying initiation. While U50 reduced the size of DA ramps overall, this did not apply to the first press of a session, which was still larger than subsequent presses in U50 trials, and there were no correlations between the size of DA ramps and the time when the lever was pressed compared to when it became available. Dopamine ramping is known to be stronger for a reliable, known reward when visual cues are present that promote anticipation ^67^. Thus, it is possible that altering a dopamine ramp through stimuli or pharmacology could affect the anticipation of and subsequent motivation to obtain a known and readily available natural reward. However, we show that rapid DA ramps were still important for task initiation during KOR activation. A more complex task like progressive ratio would be required to separate the behavior-invigorating versus motivational effects of DA ramping, and future studies are required to determine whether KOR diminishment of ramps would reduce breakpoint on a PR schedule.

Rather than affecting initiation of behavior after sucrose became available through rapid DA ramps, we found that U50 strengthened the relationship between prolonged DA ramps and subsequent responding. Of relevance to the current study, DA ramps are involved in initial learning of instrumental tasks and subsequent performance after the task is learned. For example, DA ramps in the NAc can develop within a single session ^37^, and trials preceded by increasing activity of VTA dopamine neurons with a positive slope predicted better performance in the subsequent trial compared to trials that came after a negatively sloped DA ramp ^49^. Though our paradigm was simple, with two active levers that always resulted in reward delivery, these studies align with the present results that DA activity during the 10 seconds prior to a lever press predicted when the next lever pressing bout would occur. Most interestingly, this relationship was the strongest after injection of U50, no matter how DA ramps were quantified in this 10-second pre-press period.

While the rapid, phasic-like 0.25-second DA ramp was diminished by U50, the size of prolonged DA ramps was the same between drug conditions. As KOR activation diminishes extracellular DA concentrations ^74–76^, in addition to rapid, phasic-like spontaneous DA signals ^66^, we propose that an acute hypodopaminergic state induced by U50 would increase the signal-to-noise ratio of slow, behaviorally relevant DA signals. If this occurs before a previously conditioned behavioral response, the decision to perform the action for a known outcome might be more salient, and this increased attention to the task may strengthen the relationship between DA processing and future behavior. This may be of particular importance in disorders where there is higher KOR function in the NAc, such as substance use disorders ^17,77,78^ directing behavior towards habitual, maladaptive drug responding.

This study focused on how KOR pharmacology affected operant behavior and the DA activity surrounding it with sexes combined, given that the primary effects of U50 to increase latency to respond and reduce the size of DA ramps occurred in both sexes. However, there were sex differences in overall responding, as female mice responded more and consumed more sucrose than males.^79,80^ The present results align with the literature which consistently shows that female rodents respond more robustly for and consume more sucrose than males ^79–82^ (though some studies have found that males either respond more than ^83^ or similar to females for sucrose ^84^. Studies have shown that females exhibit greater motivation to obtain sucrose, and they respond more for sucrose with increased active operant responding ^80,81^. In the present study, females responded approximately three times more for sucrose than males. Additionally, females have been shown to exhibit greater binge consumption during limited access drinking paradigms or during sucrose deprivation after a continuous schedule ^82^. This aligns with our finding that females frequently completed the maximum allowed number of responses due to more rapid lever pressing bouts with more presses per bout^79,80^.

## 5. Conclusions

The primary behavioral effect of the KOR agonist U50,488H was to delay the initiation and increase the inter-bout interval of lever pressing for sucrose, without affecting the overall number of lever presses or consumption. U50,488H diminished the rapid DA ramp that occurred immediately prior to lever pressing, though this appeared unrelated to the delay in initiation. While KOR activation did not reduce the overall DA activity 10 seconds before a lever press, it more strongly tied this activity to subsequent behavior. We propose that basal DA signaling is reduced by KOR activation, allowing a slow DA ramp to have a greater signal-to-noise ratio compared to a baseline state, thus allowing DA ramps to more precisely fine-tine future behavior under a condition of greater KOR activity. It is possible that KORs influence performance in reward-based tasks by modulating DA ramps, but more research is needed to determine whether KOR-inhibition of rapid DA ramps might be more directly tied to delays in motivated responding under more complex tasks. Overall, we report a novel effect of KOR signaling *in vivo* to reduce DA ramps, which may help in future studies in uncovering new mechanisms by which KORs mediate control over motivation and anticipation for rewards. This represents an exciting new territory to explore unknown effects of KORs to influence dopamine reward *in vivo*, including dopamine dysregulation induced by substance use disorders and effects of new pharmacological treatments on DA.

