## Supplemental data for "Kappa opioid receptors delay sucrose self-administration by modulating dopamine ramps before operant responding"

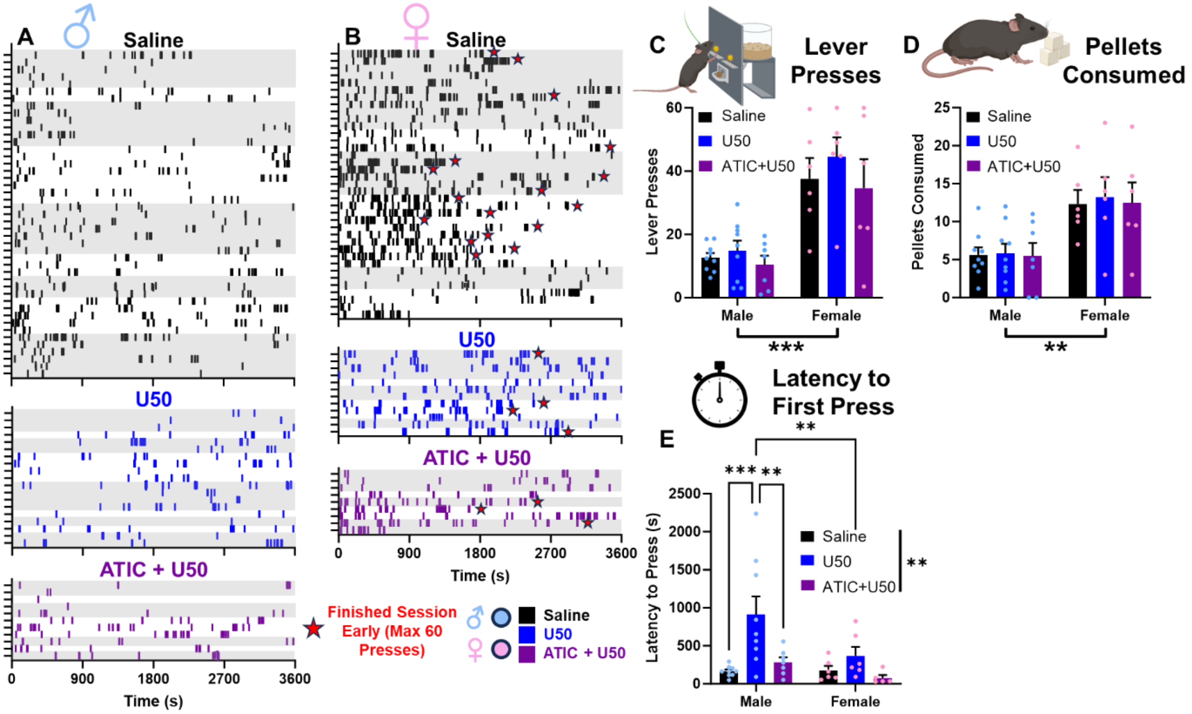

**Supplementary Figure 1. Raw behavior data.** All lever presses are represented by tick marks over hour long-sessions for **(A)** males and **(B)** females. Repeated sessions within an animal are shown within each gray or white shaded area. Only females finished sessions early (red star). There was a sex effect, but no effect of pharmacology, on **(C)** total lever presses and **(D)** pellets consumed, with females pressing and eating more. There was a significant effect of pharmacology on **(E)** latency to initiate lever pressing, with U50 delaying the start of behavior in males.

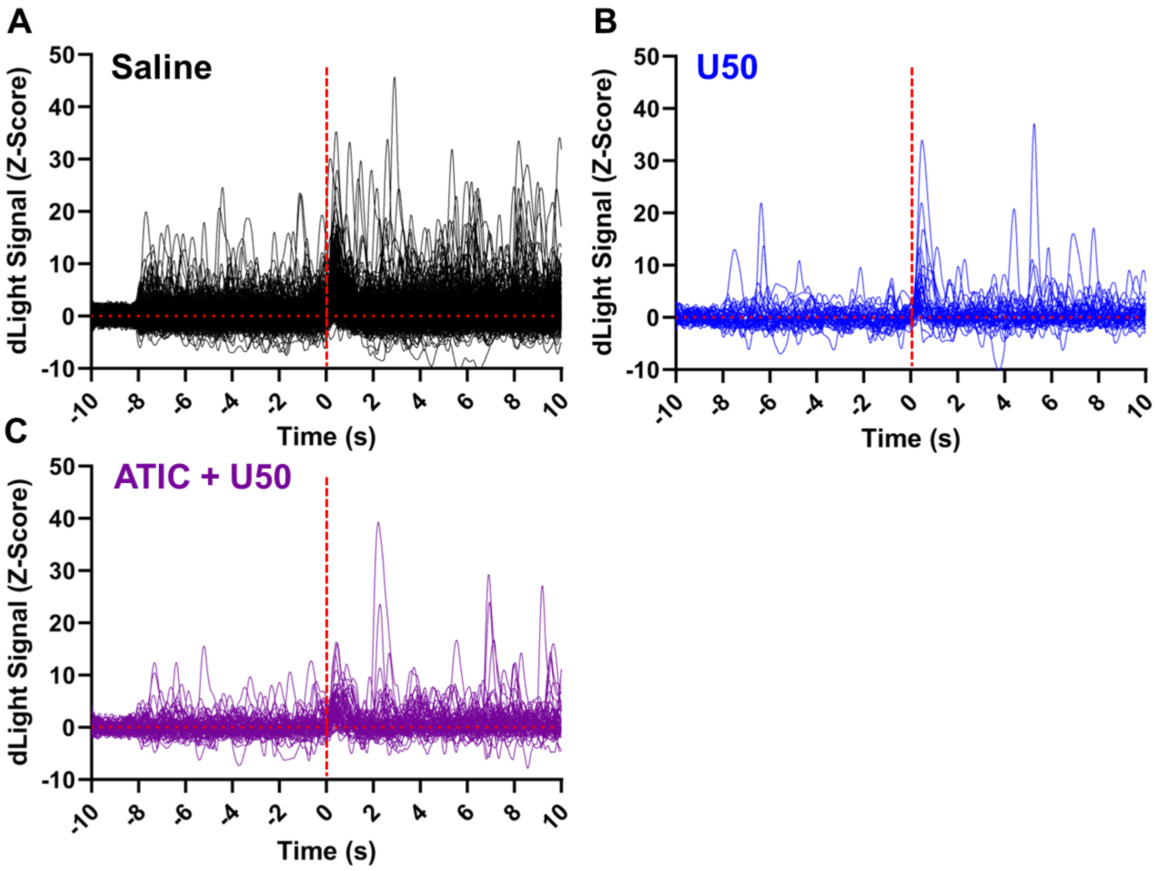

**Supplementary Figure 2. All dopamine responses to lever press bouts (-10 to 10 seconds) included in analyses. (A)** All dopamine responses (Z-Score) during saline trials. **(B)** All dopamine responses during U50 trials. **(C)** All dopamine responses during ATIC + U50 trials.

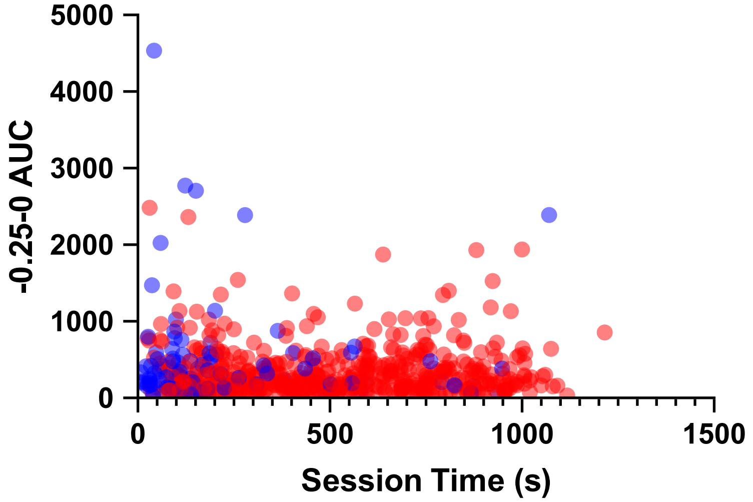

**Figure S3. Multivariable linear regression shows the first lever pressing bout in a session is preceded by a larger, rapid DA ramp.** We conducted this analysis to determine primarily whether session time or bout order related to the size of DA ramps. While those variables could not predict ramp size, DA ramps that preceded the first bout in a session tended to be larger. All responses are plotted according to ramp size by session time, with the first bout in each session highlighted.

| Model<br>Analysis of Variance | SS | DF | MS | $F_{DFn,DFd}$ | P value |
|---|---|---|---|---|---|
| Regression | 2.26E+08 | 7 | 32272759 | $F_{7,584}$ = 1.226 | 0.2864 |
| Session Time (s) | 1101805 | 1 | 1101805 | $F_{1,584}$ = 0.04184 | 0.8380 |
| Bout Order | 946168 | 1 | 946168 | $F_{1,584}$ = 0.03593 | 0.8497 |
| Presses/Bout | 2913585 | 1 | 2913585 | $F_{1,584}$ = 0.1106 | 0.7395 |
| First/Non-First | 67156420 | 1 | 67156420 | $F_{1,584}$ = 2.550 | 0.1108 |
| Sex | 96665418 | 1 | 96665418 | $F_{1,584}$ = 3.671 | 0.0559 |
| Drug | 19642630 | 2 | 9821315 | $F_{2,584}$ = 0.3730 | 0.6888 |
| Residual | 1.54E+10 | 584 | 26332694 | | |
| Total | 1.56E+10 | 591 | | | |

| Parameter estimates | Variable | Estimate | Standard error | 95% CI (asymptotic) | \|t\| | P value |
|---|---|---|---|---|---|---|
| **β0** | **Intercept** | **13312** | **637.3** | **12060 to 14563** | **20.89** | **<0.0001****** |
| β1 | Session Time (s) | 0.212 | 1.037 | -1.824 to 2.248 | 0.2046 | 0.838 |
| β2 | Bout Order | 14.95 | 78.87 | -140.0 to 169.9 | 0.1896 | 0.8497 |
| β3 | Presses/Bout | 38.63 | 116.1 | -189.5 to 266.7 | 0.3326 | 0.7395 |
| β4 | First/Non-First[Non-First] | -1030 | 645.1 | -2297 to 236.8 | 1.597 | 0.1108 |
| β5 | Sex[Female] | -985.5 | 514.4 | -1996 to 24.72 | 1.916 | 0.0559 |
| β6 | Drug[U50] | 50.85 | 867.6 | -1653 to 1755 | 0.05861 | 0.9533 |
| β7 | Drug[ATIC+U50] | -632.2 | 745 | -2095 to 831.0 | 0.8486 | 0.3965 |

**Supplementary Table 1. A multivariable linear regression analysis to predict the total AUC from -10 seconds up to the lever press from multiple behavioral and categorical variables, with interactions.** No significant independent variables were identified to predict the total AUC 10 seconds before a lever press. There was a near-significant trend for females to have reduced overall DA activity 10 seconds before the press.

| Model | | | | | |
|---|---|---|---|---|---|
| **Analysis of Variance** | **SS** | **DF** | **MS** | **F (DFn, DFd)** | **P value** |
| **Regression** | **634682** | **9** | **70520** | **F (9, 479) = 6.886** | **P<0.0001** |
| Residual | 4905173 | 479 | 10240 | | |
| Total | 5539855 | 488 | | | |

| Parameter estimates | Variable | Estimate | Standard error | 95% CI (asymptotic) | \|t\| | P value |
|---|---|---|---|---|---|---|
| **β0** | **Intercept** | **140.9** | **14.68** | **112.0 to 169.7** | **9.597** | **<0.0001****** |
| β1 | -0.25-0 AUC | 0.01242 | 0.02469 | -0.03609 to 0.06094 | 0.5032 | 0.6151 |
| **β2** | **Sex[Female]** | **-56.12** | **16.36** | **-88.27 to -23.97** | **3.430** | **0.0007***** |
| β3 | Drug[U50] | 2.981 | 49.38 | -94.04 to 100.0 | 0.06038 | 0.9519 |
| β4 | Drug[ATIC+U50] | 70.38 | 37.65 | -3.609 to 144.4 | 1.869 | 0.0622 |
| β5 | -0.25-0 AUC : Sex[Female] | -0.02637 | 0.02748 | -0.08037 to 0.02763 | 0.9596 | 0.3377 |
| β6 | -0.25-0 AUC : Drug[U50] | 0.1374 | 0.1010 | -0.06102 to 0.3358 | 1.361 | 0.1743 |
| β7 | -0.25-0 AUC : Drug[ATIC+U50] | -0.03990 | 0.05165 | -0.1414 to 0.06160 | 0.7724 | 0.4403 |
| β8 | Sex[Female] : Drug[U50] | 45.84 | 46.01 | -44.57 to 136.2 | 0.9962 | 0.3196 |
| β9 | Sex[Female] : Drug[ATIC+U50] | -39.25 | 38.67 | -115.2 to 36.72 | 1.015 | 0.3105 |

**Supplementary Table 2. A multivariable linear regression analysis to predict the time to the next bout from the size of the rapid, 0.25 second DA ramp before the lever press.** The size of the rapid, 0.25-second ramp could not predict time to the next bout, nor could any other variable other than sex, replicating findings from the other regression analyses that females had less time between bouts than males.
